# Germline and somatic variation influence nuclear-mitochondrial crosstalk in tumourigenesis

**DOI:** 10.1101/2025.02.11.637769

**Authors:** Maëlle Daunesse, Vasavi Sundaram, Elissavet Kentepozidou, John Connelly, Craig J. Anderson, Liver Cancer Evolution Consortium, Núria López-Bigas, Colin A. Semple, Duncan T. Odom, Martin S. Taylor, Maša Roller, Sarah J. Aitken, Paul Flicek

**Affiliations:** European Molecular Biology Laboratory, European Bioinformatics Institute, Wellcome Genome Campus, Hinxton, UK; Medical Research Council Toxicology Unit, University of Cambridge, Cambridge, UK; Medical Research Council Human Genetics Unit, Institute of Genetics and Cancer, University of Edinburgh, Edinburgh, UK; Edinburgh Pathology, Institute of Genetics and Cancer, University of Edinburgh, Edinburgh, UK; Laboratory Medicine, NHS Lothian, Edinburgh, UK; Institute for Research in Biomedicine (IRB Barcelona), The Barcelona Institute of Science and Technology, Barcelona, Spain; Universitat Pompeu Fabra (UPF), Barcelona, Spain; Institució Catalana de Recerca i Estudis Avançats (ICREA), Barcelona, Spain; Division of Regulatory Genomics and Cancer Evolution (B270), German Cancer Research Center (DKFZ), Heidelberg, Germany; Cancer Research UK - Cambridge Institute, University of Cambridge, Cambridge, UK; Center of Molecular and Cellular Oncology, Yale Cancer Center, New Haven, USA; Department of Pathology, Yale School of Medicine, New Haven, USA; Department of Histopathology, Cambridge University Hospitals NHS Foundation Trust, Cambridge, UK; The Jackson Laboratory for Genomic Medicine, Farmington, USA; Department of Genetics, University of Cambridge, Cambridge, UK

## Abstract

Hepatocellular carcinoma is the most prevalent primary liver cancer and arises from hepatocytes, which are very metabolically active and contain abundant mitochondria. Despite extensive characterisation of nuclear driver genes and pathways, the role of crosstalk between the mitochondrial and nuclear genomes in mitochondrial mutagenesis remains poorly understood. To address this, we leveraged a chemical carcinogenesis model of liver cancer in four mouse strains. We analysed hundreds of tumours with paired whole-genome sequencing and total RNA sequencing to comprehensively characterise mitochondrial mutations and expression. Following *de novo* assembly of strain-specific mitochondrial genomes, we devised an accurate and efficient heteroplasmy detection approach and analysed mitochondrial DNA (mtDNA) content and transcription. Across strains, there was high concordance of the number of heteroplasmies, mutation signatures, and variant allele frequencies. In contrast, the heteroplasmy rate and genic locations differed between strains, suggesting strain-specificity in DNA repair mechanisms in early tumour development. Independent of age or environmental factors, tumours had lower mtDNA content than adjacent normal liver, suggesting mitochondrial loss during tumourigenesis. Additionally, within individual strains, mtDNA content differences were associated with nuclear driver gene choice. Our comprehensive characterisation of mitochondrial genomes and expression patterns demonstrates that both germline and somatic genetic variation influence tumour mtDNA content, mutation patterns, and gene expression.

## Background

Hepatocellular carcinoma (HCC) is the most common type of primary liver cancer. It has the sixth highest incidence of all human cancers with five-year survival of only 12% (Bray et al. 2018). HCC typically occurs in the context of chronic liver disease (Llovet and Beaugrand 2003) caused by repeated damaging exposures, such as viral hepatitis infection, alcohol consumption, or metabolic dysfunction-associated steatotic liver disease (MASLD). Although there are considerable differences in HCC burden across the globe (Toh et al. 2023), how an individual’s genetic variation interacts with these known environmental risk factors is largely unstudied.

Cancer initiation and promotion, including that of HCC, is the product of acquired somatic mutations (Stratton et al. 2009). To identify such mutations within nuclear DNA, whole exome (Tomczak et al. 2015) and whole genome (Aaltonen et al. 2020; Zhang et al. 2019a) sequencing has been applied to large cohorts of liver cancers. These efforts have led to the identification of multiple driver gene mutations in distinct oncogenic pathways involved in HCC development (Schulze et al. 2015; Zucman-Rossi et al. 2015; Fujimoto et al. 2016; Ng et al. 2021; Wu et al. 2023), as well a better understanding of somatic mosaicism in chronic liver disease (Brunner et al. 2019; Ng et al. 2021). Despite this extensive characterisation of the nuclear genome, the contribution of mutations in the mitochondrial genome to liver tumourigenesis remains understudied.

Mitochondria are eukaryotic cytoplasmic organelles colloquially known as the “powerhouses of the cell”. Mitochondria synthesise adenosine triphosphate (ATP) through oxidative phosphorylation to provide the energy required to support cellular functions (Osellame 2012). Mammalian cells generally each have between 100 and 10,000 mitochondria depending on their function and developmental stage. Each mitochondrion contains one or more copies of its own extra-nuclear circular genome, referred to as mitochondrial DNA (mtDNA). In mice, as in humans, the mtDNA is approximately 16 kb in size and it contains 37 genes encoding two rRNAs, 22 tRNAs, and 13 protein-coding genes that are components of the oxidative phosphorylation system. Mutations in mtDNA are mostly single nucleotide substitutions, insertions, or deletions (Kogelnik et al. 1996), typically occurring during mitochondrial genome replication and persisting in the mitochondrial population through repeated mitotic segregation. These mutational events can lead to multiple mtDNA variants in a cell or individual: A phenomenon known as heteroplasmy.

Liver cells (hepatocytes) have very high ATP requirements to perform their multiple energy-consuming functions, including detoxification of blood and the metabolism of fat, proteins, and carbohydrates. Indeed, high-resolution imaging of mouse hepatocytes estimates the number of mitochondria to be between 1,000 and 3,000 per cell (Parlakgül et al. 2024), making the liver tissue one of the most abundant sources of mtDNA. Deregulation of cellular metabolism is recognised as a core hallmark of cancer (Hanahan and Weinberg 2011; Hanahan 2022). More specifically, mitochondria are thought to play a central role in cancer development because malignant cells are characterised by increased cell proliferation, resistance to cell death, and high energy consumption (Hertweck and Dasgupta 2017). Recent studies have suggested that disruptive mutations to mtDNA genes act as drivers in particular tumour types (Kim et al. 2022), and more generally, in normal tissues, both mtDNA copy number and heteroplasmy behaviour have been shown to be under nuclear genetic control in a large human genome wide association study (Gupta et al. 2023).

However, we still have a limited understanding of the mutagenic processes that occur in mtDNA during tumourigenesis and the functional impact of these mutations.

Given the substantial energy requirements of liver tissue and importance of metabolic alterations in HCC (Yang et al. 2023), mtDNA is a critical component of liver cancer genome analysis. Indeed, previous work has identified global heteroplasmy patterns, expression changes, and other dysregulation in mtDNA associated with human liver tumourigenesis (Reznik et al. 2016; Yuan et al. 2020). Currently, the functional consequence of those heteroplasmies remains unclear; some studies showing positive selection for somatic mutations (Li et al. 2015; Grandhi et al. 2017) while others found neutral evolution (Liu et al. 2019; Yuan et al. 2020). There is also evidence for decreased mitochondrial copy number (Reznik et al. 2016; Yuan et al. 2020) and decreased expression of mitochondrial genes (Reznik et al. 2017; Yuan et al. 2020) in human liver cancer compared to the tissue and cells of origin. In other tumour types, mtDNA mutation load has been identified as a potential biomarker of treatment response (Mahmood et al. 2024), as well both a positive (Gorelick et al. 2021) and negative (Ewing et al. 2024) overall survival in colorectal and ovarian cancer, respectively. However, these studies did not consider individuals’ genetic background.

To investigate how individual genetic background affects mitochondria during tumourigenesis we used a well-characterised experimental model of liver cancer and analysed nearly six hundred liver tumours spanning four genetically and evolutionary distinct mouse strains (Connor et al. 2018; Aitken et al. 2020, 2025). By controlling for both genetics and environment, we could determine the consequences of differences in genetic background on mtDNA mutations, content, and expression.

## Results

We chemically induced liver tumours by treating 15-day-old mice with a single dose of diethylnitrosamine (DEN), and performed whole genome sequencing (WGS), total transcriptome sequencing (RNA-seq), and histopathological analysis (Connor et al. 2018; Aitken et al. 2020, 2025). For each strain, tumours were collected at a defined time point post DEN treatment and histologically confirmed as dysplastic nodules (DNs). For this study, we included tumours (n=580) from inbred laboratory *Mus musculus* C3H/HeOuJ (C3H) mice (n=370 DNs) and C57BL/6J (BL6) mice (n=55 DNs); wild-derived inbred *Mus musculus castaneus* (CAST) mice (n=84 DNs); and wild-derived inbred *Mus caroli* (CAROLI) mice (n=71 DNs) (**Table S1**). The majority of the CAST genome separated from the common ancestor of BL6 and C3H approximately 0.5 million years ago, although there are introgressions of the ancestral CAST genome in the BL6 and C3H genomes (Yang et al. 2011). CAROLI diverged from the common ancestor of the other three strains approximately 3 million years ago (Thybert et al. 2018). For clarity, these species, subspecies, and strains are all subsequently referred to as strains. We have recently shown that these germline differences influence cancer susceptibility and somatic mutagenesis in the nuclear genome (Aitken et al. 2025). These strain-specific differences include driver gene preference within the MAPK pathway, genome stability (including whole genome duplication), and tumour latency which ranges from 25 weeks for C3H to 80 weeks for CAROLI.

### *De novo* assembly of novel mitochondrial genomes

To ensure the robustness of our cross-strain analyses, we needed to use strain-matched mitochondrial genomes. Therefore, we assembled and annotated de novo mitochondrial genomes using our WGS data generated from matched non-tumour samples (Aitken et al. 2020, 2025) from C3H, CAST, and CAROLI (Methods) and used these mitochondrial genomes in conjunction with the annotated reference BL6 mitochondrial genome (NC_005089.1) (Bayona-Bafaluy et al. 2003). Our comparative analysis confirmed there are no structural rearrangements between these four mitochondrial genomes, they contain the same set of 37 genes (**Table S2**), and they are of comparable genomic size (C3H 16,301 bp; BL6 16,299 bp; CAST 16,299 bp; CAROLI 16,310 bp) (**Fig. 1**). Additionally, the sequence divergence between genomes reflects the evolutionary distance of the four strains (**Fig. S1a**): Specifically, closely related BL6 and C3H are 99.98% identical, whereas more divergent strains CAST and CAROLI exhibit lower sequence identity to BL6, at 97.67% and 87.21%, respectively (**Fig. S1b**).

**Figure 1.**
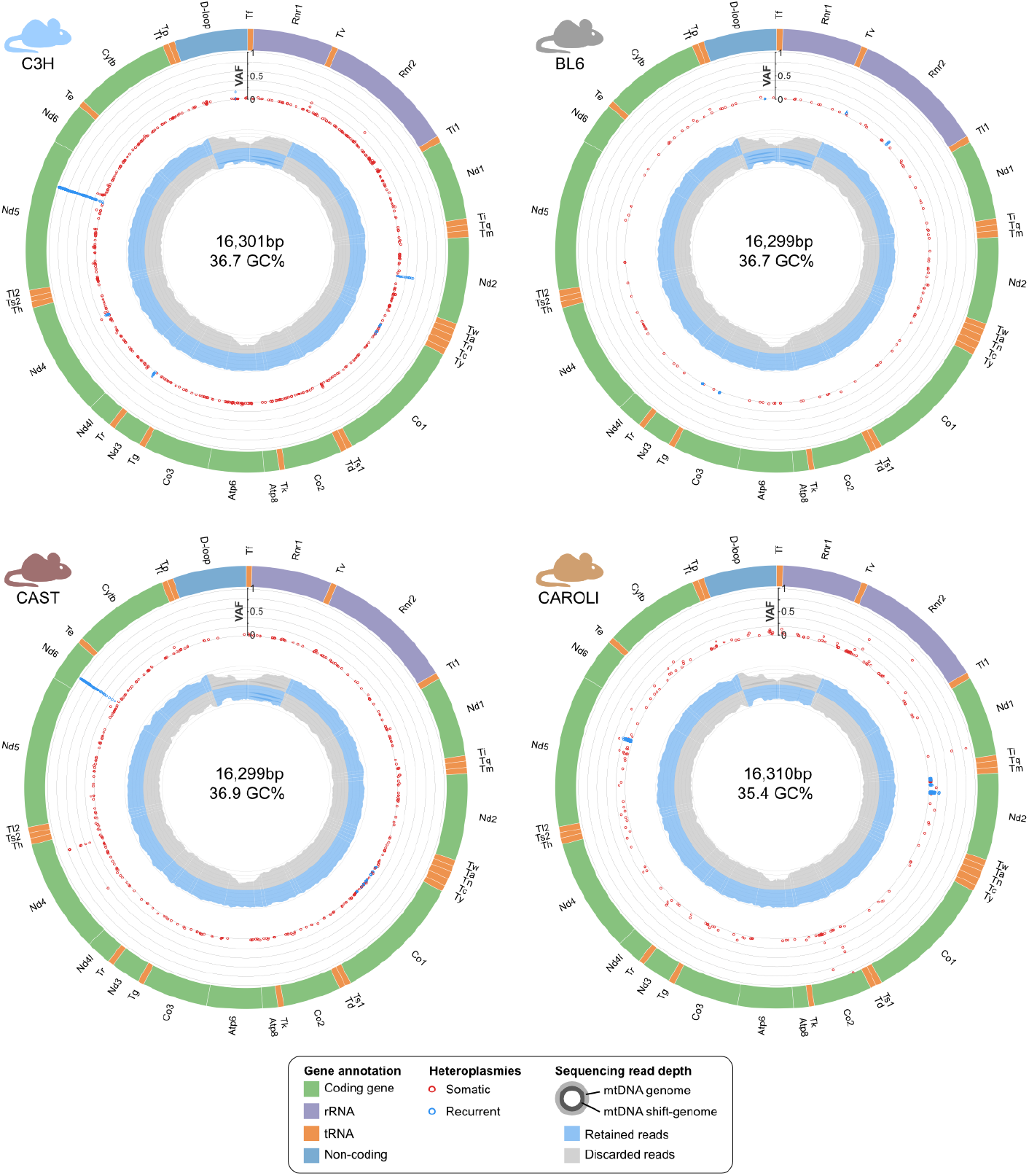
Mitochondrial genome assemblies demonstrate similar genome structure, size, GC content, and sequencing coverage across strains. The C3H, BL6, CAST, and CAROLI mitochondrial genomes represented in CIRCOS plots. The outer layer of each plot represents the mitochondrial genomic position with gene annotation. The intermediate layer shows heteroplasmy variant allele frequency for somatic mutations (red) and recurrent heteroplasmies (blue) calibrated on the y-axis. The inner layer shows mitochondrial sequencing coverage for normal (external) and shift (internal) mapping (see **Fig. S3)**. The genomic regions that are retained from each mapping approach are shaded blue (Methods).

To validate the accuracy of our *de novo* assemblies, we compared our assemblies to other available mtDNA assemblies (**Table 1**; Methods). Our genome assembly for CAST is 99.99% identical to the reference CAST mtDNA (NC_012387.1) with only two substitutions (positions 2,797 and 13,733). For CAROLI, our assembly is 98.88% identical to the reference *Mus caroli* mtDNA (NC_025268.1) and 99.99% identical to another *Mus caroli* mtDNA (MK166027.1). This difference likely arises from the fact that both our samples and those used for MK166027.1 were obtained from the same commercial provider, whereas the reference *Mus caroli* mitochondrial genome (NC_025268.1) was sequenced from a wild-caught individual (Rachal O’Neill, Johan Michaux, and Alex Greenwood personal communication). Although we were not able to identify a previous mtDNA assembly for our C3H strain (C3H/HeOuJ), comparison to a C3H/HeJ mitochondrial genome (EF108335.1) showed 99.98% identity, with only three substitutions (positions 8,898, 9,821, and 13,066). Taken together, these comparisons confirm that our *de novo* mitochondrial genome assemblies are high accuracy and consistent with published references, while also revealing minor sequence variations likely reflecting natural genetic drift.

**Table 1.** Mitochondrial genomes included in the comparative analysis.

| Strain | GenBank ID | Length (bp) | Strain information |
| --- | --- | --- | --- |
| **C3H** | **PV177089.1** | **16,301** | ***Mus musculus* strain C3H/HeOuJ (this study)** |
| C3H | EF108335.1 | 16,301 | *Mus musculus* strain C3H/HeJ |
| **CAST** | **PV231059.1** | **16,299** | ***Mus musculus castaneus* strain CAST/EiJ (this study)** |
| CAST | EF108342.1 | 16,299 | *Mus musculus castaneus* strain CAST/EiJ<br>Curated RefSeq NC_012387.1 |
| **CAROLI** | **PV231060.1** | **16,310** | ***Mus caroli* strain CAROLI/EiJ (this study)** |
| CAROLI | KJ530558.1 | 16,313 | *Mus caroli* isolate R5231 mitochondrion<br>Curated RefSeq NC_025268.1 |
| CAROLI | MK166027.1 | 16,301 | *Mus caroli* |

We next examined the sequence conservation of the origin of replication for the light strand (ORI-L), an essential region for mtDNA maintenance in mammals (Wanrooij et al. 2012), which contains a highly conserved polyA sequence whose length has been associated with lifespan in mice (Hirose et al. 2018). In C3H, BL6, and CAST the ORI- L sequence is fully conserved, including an 11-adenine repeat (**Fig. S2**). However, in CAROLI, this region has evolved into a 10AG3A sequence, including a G-to-A change (**Fig. S2**). These differences at the CAROLI ORI-L locus are of particular interest given the longer lifespan and increased cancer resistance observed in CAROLI mice.

### Novel heteroplasmy detection workflow using whole genome sequencing

To detect heteroplasmies with high confidence across the entire mitochondrial genome, we implemented a novel heteroplasmy detection workflow, taking the circular nature of the mitochondrial genome into consideration. Our mapping approach (**Fig. S3a**) builds on a previous study (Zhang et al. 2019b), with a double alignment step (**Fig. S3b**), which reduces the detection of false positive somatic heteroplasmies, particularly those caused by the misalignment of mtDNA fragments integrated into the nuclear genome termed nuclear mitochondrial sequences (NUMTs).

First, we mapped whole genome sequencing reads to the mitochondrial genome. Uniquely mapping, valid, paired reads were then extracted and re-mapped to both the nuclear and mitochondrial genomes; only those that aligned to the mtDNA (and not to the nuclear genome) were extracted, thus ensuring that only true mitochondrial reads were considered for the next steps. These reads were then deduplicated and homoplasies were called and annotated (Methods; **Fig. S3a**), with an average sequencing depth of 5,166X at heteroplasmic loci. To account for the circularity of the mitochondrial genome, we created an alternate version of the mtDNA assembly for each of the four strains by shifting the linear starting position of the mtDNA by 8,150 bp (**Fig. S3b**). Then, the mapping approach described above was reapplied to this shifted genome, which results in more even read depth across the entire mitochondrial genome, in particular at the start and end of the mtDNA coordinate system which otherwise suffer from lower coverage due to unpaired read alignment (**Fig. 1**). A final heteroplasmy set was obtained by combining the heteroplasmies identified using the reference genome (nucleotide position 1,100 to 15,200) and the shifted genome (position 1 to 1,099, and 15,201 to the end of the genome) (**Fig. 1**; **Fig. S3b**).

Our novel approach of repeating the mapping using a shifted mtDNA reference (in addition to the classic reference) improved the coverage by an average of 1,500X in regions near the start and end of the mtDNA genome (**Fig. S5**). In addition to ensuring more uniform mapping coverage, this approach also significantly improves accuracy and specificity: In this dataset, we were able to exclude 310 false positive heteroplasmies that would otherwise be erroneously detected solely because of low coverage.

### Strain-specific differences in mutational landscapes of tumour mitochondrial genomes

To identify and characterise the mitochondrial mutation landscape, we performed heteroplasmy detection on all normal (n=27) and tumour (n=580) samples for each of the four strains (Methods). Since germline selection can shape mtDNA diversity in humans (Wei et al. 2019), we first analysed the genomes of normal liver samples to identify germline variation/polymorphisms in each strain (**Fig. 1**). On average, each normal liver tissue genome had one heteroplasmy; since normal tissue is expected to be polyclonal, these heteroplasmies were considered to be germline in origin and therefore removed from the tumour set. Additionally, we identified that 49% of the heteroplasmy positions in tumours are strain-recurrent (Methods). The variant annotation of the strain-recurrent heteroplasmies revealed that 78% are synonymous and none were annotated as loss of function. Because these recurrent heteroplasmies are likely to be germline or NUMT false positives, we also excluded them from downstream analysis.

After removing hypermutated tumours (n=7; Methods; **Fig. S6**), the remaining 573 tumour samples (C3H n=369; BL6 n=55; CAST n=80; CAROLI n=69) contained a total of 1,138 somatic heteroplasmies (C3H n=527; BL6 n=126; CAST n=340; CAROLI n=145; **Table S4**), equating to a median of 1 to 4 (C3H median=1, BL6 and CAROLI 2, CAST 4) heteroplasmic sites per genome (**Fig. 2a**). Tumour samples are significantly more mutated than normal liver (p<0.001, Wilcoxon test). Heteroplasmies are positively correlated with nuclear somatic mutation rate in each strain (**Fig. S6**); accordingly, BL6 and CAST tumours have more mitochondrial heteroplasmies than those from C3H and CAROLI, also reflecting the elevated levels of mutation in the nuclear DNA for these strains (Aitken et al. 2025). The mutational spectra signature is nearly identical across the four strains. The majority of mutations are C:G>T:A substitutions (∼60%) and T:A>C:G substitutions (∼30%) (**Fig. 2b**). This mutation profile does not reflect the DEN signatures that dominate the tumour nuclear genomes (Aitken et al. 2025); instead, it closely matches the mtDNA signatures seen in human cancers (Lake et al. 2024; Yuan et al. 2020) which reportedly reflects mtDNA replication strand biases (Ju et al. 2014).

**Figure 2.**
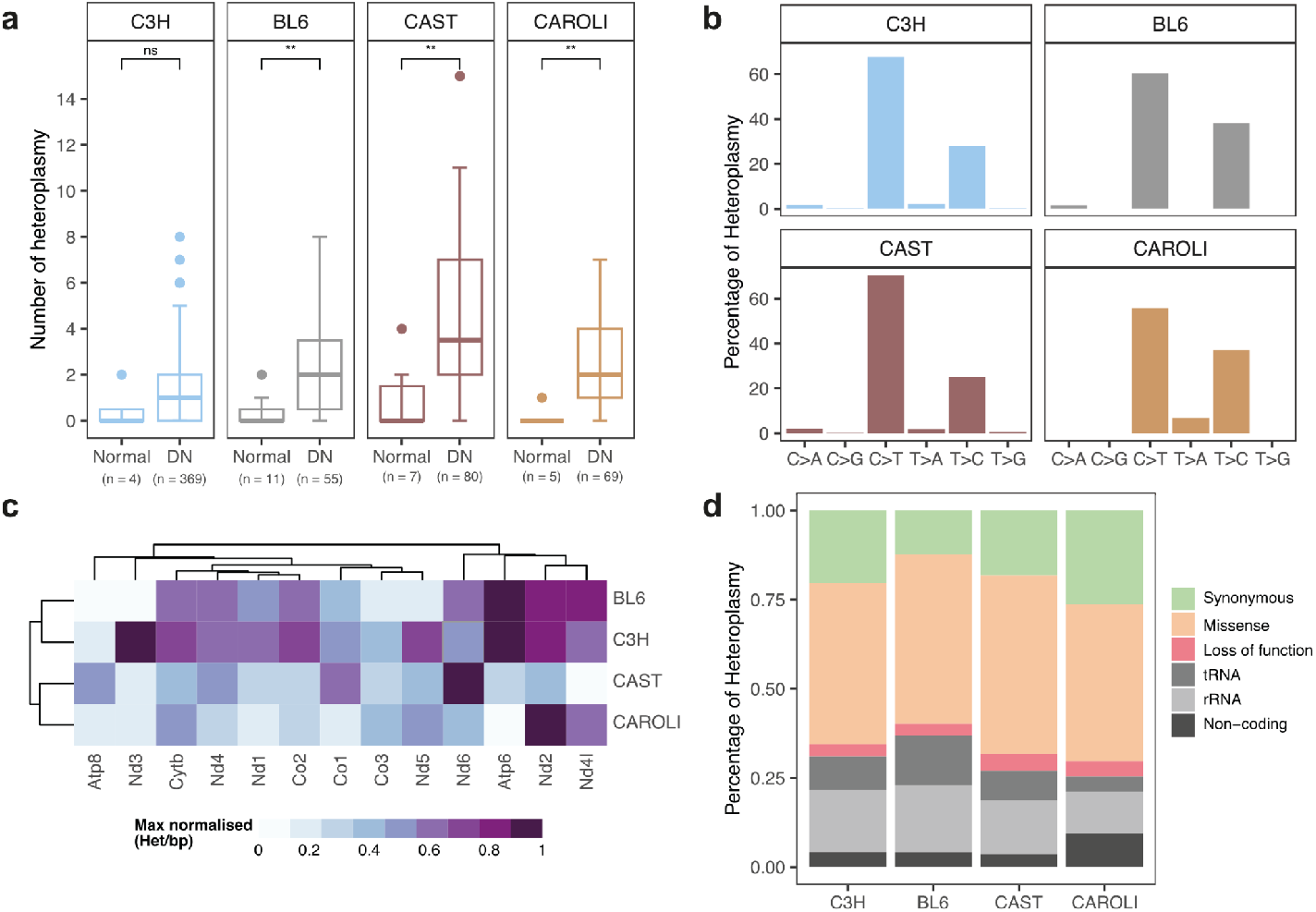
Characterisation of heteroplasmies in normal liver and tumour samples. **(a)** Liver tumour (DN) genomes contain more mitochondrial heteroplasmies than normal liver tissue, which is replicated across all strains (C3H, BL6, CAST, CAROLI). P-values were calculated using a Wilcoxon test (ns = not significant, p>0.05; ** = p<0.01). **(b)** Percentage of heteroplasmies associated with each nucleotide substitution type (C>A, C>G, C>T, T>A, T>C, T>G) was calculated across all tumour samples on a per-strain basis; percentages include reverse complement mutations (e.g. T>C includes A>G). **(c)** Sequence-composition adjusted mutation rate (max normalised heteroplasmy per base pair) per gene, normalised by strain. The heatmap displays the heteroplasmy rates across mitochondrial genes, with darker shades indicating a higher mutation rate. **(d)** For each of the 1,138 somatic heteroplasmies (C3H n=527; BL6 n=126; CAST n=340; CAROLI n=145), the variant type was annotated (synonymous, missense, loss of function, tRNA, rRNA, and non-coding) and the proportions calculated per strain. Approximately 40% of variants are non-synonymous and may impact protein function.

The majority of heteroplasmies (75%) have a low variant allele frequency (VAF; <2%), which is consistent with the accumulation of replication-coupled mutations (Ju et al. 2014). Given the gradual acquisition of such mutations, and the observation in humans that heteroplasmic single nucleotide variants can increase with age (Gupta et al. 2023), we considered whether this also applies in mice. We detected only a weak positive correlation between the number of heteroplasmies per sample and the age of the mouse at tumour isolation (Spearman’s ρ=0.29, p<0.001; **Fig. S7a**). Similarly, VAF is only very weakly correlated with age (Spearman’s ρ=0.08, p<0.001; **Fig. S7b**).The absence of a stronger correlation in our cohort may arise because the experimental design imposed a fixed time point per strain (Aitken et al. 2025), which means that age is fully confounded with genetic background.

Next, we focussed specifically on mutations arising within the thirteen mitochondrial protein-coding genes. We calculated sequence-composition adjusted mutation rate for each gene, which showed heterogeneity among the four strains (**Fig. 2c**). We find no significant difference between the genic distribution of heteroplasmies in C3H and BL6 (p=0.22), most of which are within the NADH dehydrogenase complex. Heteroplasmies in the other strains are less evenly distributed, with mitochondrially encoded NADH dehydrogenase 6 (*Nd6*) overrepresented in CAST, and mitochondrially encoded NADH dehydrogenase 2 (*Nd2*) and mitochondrially encoded NADH dehydrogenase 4L (*Nd4l*) in CAROLI. The NADH dehydrogenase complex accounts for 39% of the mitochondrial genome, and somatic mutations have been associated with clinical outcome in diverse human cancers (Kurelac et al. 2013; Koshikawa et al. 2017).

To understand the impact of these protein coding mutations we annotated their functional consequences and found a high proportion of missense mutations (∼40%) in all four strains (**Fig. 2d**). This is consistent with previous work which found that, in contrast with the nuclear genome, mtDNA shows an enrichment for non-synonymous variants, including in human liver cancer (Hertweck and Dasgupta 2017; Yuan et al. 2020). We found that CAST has a significantly higher proportion of heteroplasmies with a loss of function annotation compared with the three other strains (Fisher’s exact test p=0.033) (**Fig. 2d**).

We measured the relative rates of non-synonymous (dN) to synonymous (dS) mutations for each mitochondrial gene to infer selective pressure (**Table S3**). The resulting mtDNA-wide dN/dS scores suggest neutral evolution for missense heteroplasmies in C3H (1.0) and CAST (1.0). However, BL6 shows evidence of positive selection for missense heteroplasmies with nearly twice as many observed as expected by chance (dN/dS ratio 1.9; 95% confidence interval (CI) 1.01-3.57). BL6 genes with significant evidence for positive selection include *Nd2* (dN/dS ratio 4.7) and *Nd4* (dN/dS ratio 2.7). In contrast, the CAROLI mtDNA-wide score of 0.4 (95% CI 0.27- 0.66) indicates higher purifying selection for mitochondrial mutations in this strain, with significant evidence for selection on *Nd2* (dN/dS ratio 0.2). We also estimated which of the non-synonymous heteroplasmies are likely to be deleterious by calculating SIFT scores for each of the non-synonymous heteroplasmies (Kumar et al. 2009) (**Table S4**; Methods). For each strain, approximately three quarters of the missense heteroplasmies are predicted to be deleterious.

In summary, the concordance of the number of heteroplasmies, mutation signatures, and VAF distribution across all tumours strongly suggests that the mutational mechanisms of tumourigenesis are highly conserved across all strains in this study. However, the differences in heteroplasmy rate and genic location imply that strain-specific repair mechanisms or other differences in the dynamic process of tumourigenesis play an important role. These results illustrate the importance of background genotype on patterns of somatic mutations.

### Mitochondrial content is associated with tumour development and driver gene mutations

Having defined the mutational landscape of mtDNA mutations in DEN-induced tumours, we calculated changes in mtDNA content between normal liver and tumour sample (Methods, **Table S5**). We found that tumours have significantly lower mtDNA content than normal tissue in BL6 and CAST (**Fig. 3a**). C3H tumours show the same trend, but have too few normal samples to assess statistical significance, while CAROLI tumours demonstrate a different phenomenon described below. To determine whether the observed depletion in mitochondrial content is associated with (i) DEN-treatment, (ii) ageing (the normal samples used in **Fig. 3a** are 15 days old), or (iii) tumourigenesis, we analysed two other BL6 sample sets. First, we collected background (non-tumour) liver tissue that was spatially distant to the DEN-induced tumours; by definition, these background tissues were also exposed to DEN when the mouse was 15 days old. Comparing paired DEN-exposed background liver to DEN-induced tumour from the same individuals (**Fig. 3b**), shows that tumours have significantly lower mtDNA content than the DEN-treated background samples (p<0.0001, Wilcoxon test). Furthermore, there is no significant difference in mitochondrial content between these DEN-exposed background liver samples and unexposed 15-day-old normal liver (**Fig. S7c**). This indicates that tumour development itself, rather than DEN treatment or age, is responsible for mitochondrial content loss. Second, we included a cohort of DEN-induced HCCs, which are histologically more advanced than the majority of dysplastic nodule tumours analysed in the rest of this study; this allowed us to investigate the effect of tumour stage on mtDNA content. We found that the mtDNA content of the HCC samples is comparable to the DNs (**Fig. 3b**), suggesting that while tumour development is associated with a decrease in mtDNA content, it is independent of tumour stage. These data are strikingly similar to the lower mtDNA content seen in human HCC relative to control liver tissue (Yuan et al. 2020), suggesting our murine model system recapitulates features of human disease.

**Figure 3.**
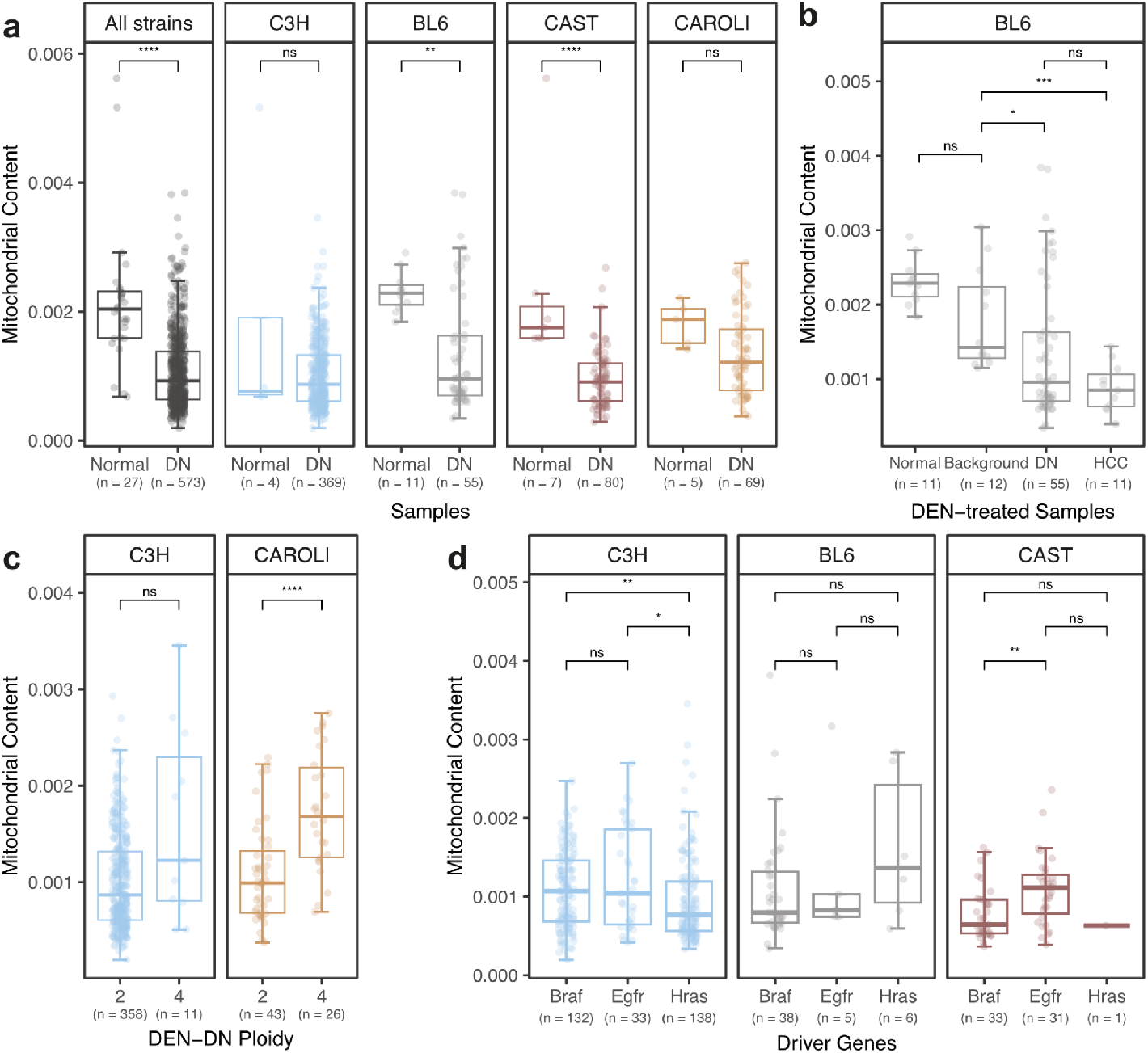
Tumourigenesis alters mitochondrial content. **(a)** Across all samples, tumours (DN) have less mtDNA content than normal liver samples, which is also significant in strain-specific analysis of BL6 and CAST. **(b)** Tumour (DN and HCC) genomes have significantly lower mtDNA content than adjacent, age-matched background liver tissue (DEN-exposed, non-tumour) from the same individual BL6 mouse, or normal (15-day-old, DEN-naive) liver, which indicates that it is the process of tumourigenesis itself, rather than DEN treatment or tumour stage, that affects mtDNA content. **(c)** Whole genome duplicated tumours (4n) have significantly increased mtDNA content compared to diploid tumours (2n) in CAROLI, consistent with both increased nuclear and cytoplasmic volume. **(d)** Mitochondrial content is affected by an interaction of genetic background and somatic mutations: C3H tumours with *Hras* mutations are significantly depleted in mitochondrial content compared with those with *Braf or Egfr* mutations, while CAST tumours with *Braf* mutations are depleted compared with *Egfr* driven tumours. For all plots p-values (p) were calculated using a Wilcoxon test (ns = p>0.05, * = p<0.05, ** = p<0.01, *** = p<0.001, **** = p<0.0001).

A distinct feature of the CAROLI cohort is the high prevalence of whole-genome duplication (WGD) (Aitken et al. 2025), which was observed in 26 out of 69 CAROLI tumours. Analysing the mitochondrial content of these WGD tumours also revealed an approximate doubling of mtDNA content compared to diploid CAROLI tumours (**Fig. 3c**). In contrast, the small proportion of whole genome duplicated C3H tumours (11/369) do not have significantly higher mtDNA content than diploid tumours (**Fig. 3c**). The CAROLI results are consistent with a failure of karyokinesis, including preservation of both pairs of sister chromatids in the same nucleus (i.e. a single, 4n nucleus per cell) and increased cytoplasmic volume, thus maintaining the nuclear to mtDNA ratio. This hypothesis is supported by image analysis of digitised histopathology which found an approximate doubling of nuclear volume in CAROLI WGD tumours (Aitken et al. 2025), and shows strong parallels with human tumours, where WGD has been shown to evolve in concert with increased mitochondrial density (Kim et al. 2024).

Finally, we considered whether specific driver gene mutations in the nuclear genome had an impact on mtDNA content. This question is effectively intractable in human cohorts where there is often high diversity and multiple combinations of driver mutations present. In our study, the vast majority of tumours across all strains have an activating mutation in the MAPK pathway, but the genic distribution of these varies between strains. For that reason, we compared driver genes on a per strain basis and focused on tumours with one of the three most prevalent driver genes (Braf transforming gene (*Braf*), epidermal growth factor receptor (*Egfr*), and Harvey rat sarcoma virus oncogene (*Hras*)); CAROLI was omitted because there were too few diploid tumours to test statistical significance. In C3H, we find significantly higher mtDNA content in *Braf* driven tumours compared to *Hras* tumours (p< 0.01, Wilcoxon test) (**Fig. 3d**). By contrast, *Egfr* driven tumours had the highest mtDNA content in CAST tumours (p<0.05, Wilcoxon test). To our knowledge, this is the first demonstration of associations between the activation of particular driver genes and mtDNA content in tumours, though it has been established that MAPK signalling can affect many aspects of mitochondrial metabolism and biogenesis (Sarg et al. 2024).

In summary, mitochondrial content (i) decreases during tumourigenesis, (ii) is associated with specific driver gene mutations, and (iii) is correlated with whole genome duplication in some genetic backgrounds.

### Tumourigenesis results in strain-specific mitochondrial expression profiles

To characterise the functional impact of mutations in the mitochondrial genome on cancer development we analysed the gene expression of all tumours with paired WGS and RNA-seq (n=378) and 52 normal liver transcriptomes (Methods). We first examined the overall expression of all thirteen mitochondrial genes (**Fig. 4a**) in normal and tumour samples across the four strains (**Table S6**). This revealed that C3H, CAST, and CAROLI have a significant loss of gene expression in tumour samples compared with normal samples, whereas BL6 tumours showed more muted changes. While the overall reduced mitochondrial gene expression is consistent with the trend for depletion of tumour mtDNA content (**Fig. 3a**), for a given tumour the correlation between mtDNA content and gene expression is relatively low for diploid tumours, with the exception of CAROLI (Spearman’s ρ=0.42; p<0.01) (**Fig. S8**). WGD CAROLI tumours show stronger correlation (Spearman’s ρ=0.57; p<0.01) than their diploid counterparts, which is also observed in the minority group of C3H WGD tumours (Spearman’s ρ=0.43; p<0.01) (**Fig. S8**).

**Figure 4.**
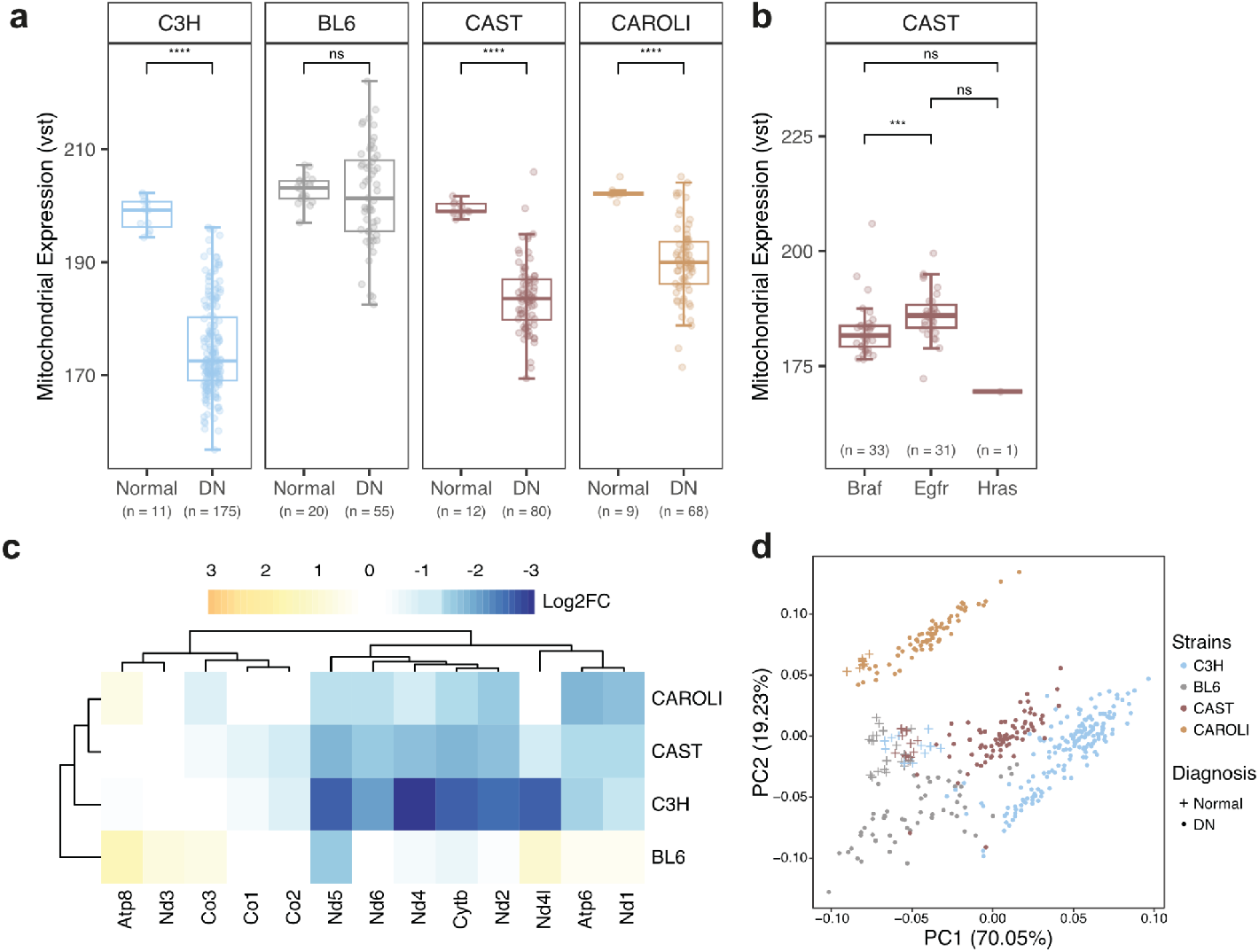
Tumour samples exhibit loss of mitochondrial gene expression. **(a)** Liver tumour (DN) samples have significantly lower mitochondrial gene expression than normal age-matched liver tissue in C3H, CAST, and CAROLI mice (Wilcoxon test, ns = p>0.05, *** = p<0.001, **** = p<0.0001; VST = variance stabilisation transformation). **(b)** In CAST mice, the expression level in tumour samples is associated with driver gene mutations, with *Egfr* mutated tumours having the highest expression levels. There was no correlation in other strains (**Fig. S9**). **(c)** Differential gene expression of mitochondrial genes was calculated between normal adult and tumour samples, per strain. As for total gene expression, the per-gene analysis highlights more marked changes in C3H, CAST, and CAROLI (Log2FC = log2 fold-change; p<0.01). Colour intensity indicates the magnitude of change. **(d)** Principal component analysis (PCA) of the VST mitochondrial expression read counts for all normal and tumour samples. Normal samples (crosses) cluster together, whereas tumour samples (circles) separate by strain.

We also investigated the potential impact of specific driver genes on mitochondrial expression. Again, we focused our comparison on tumours that have one the three most prevalent oncogenes (*Braf, Egfr*, and *Hras*) and CAROLI samples were excluded due to insufficient tumour numbers. We found that CAST *Egfr* driven tumours show significantly higher mitochondrial expression than those with *Braf* drivers (**Fig. 4b**), consistent with the higher mtDNA content observed in CAST tumours (**Fig. 3d**).

To assess the mitochondrial gene expression changes in tumours, we performed a differential expression analysis of all 378 tumour RNA-seq libraries compared to normal adult liver samples (**Fig. 4c**). The C3H, CAST, and CAROLI tumours showed lower expression in tumours for most mitochondrial genes, especially those in the NADH complex, mitochondrially encoded ATP synthase 6 (*Atp6)*, and mitochondrially encoded cytochrome b (*Cytb)*. As with overall gene expression, above, BL6 shows a blunted pattern with fewer, and less significantly, differentially expressed genes.

Finally, we performed principal component analysis (PCA) using normalised mitochondrial expression of all normal and tumour samples for the four strains (**Fig. 4d**). Most striking is the tight clustering together of C3H, BL6, and CAST normal liver samples, with slight separation by age (**Fig. S10**); this is in contrast to the tumours which diverge between strains, but cluster within strain, partially accounting for PC1. The CAROLI samples behave differently, with normal and tumour samples clustering away from the other strains (accounting for PC2). The mitochondrial genes analysed here show a profile consistent with our transcriptome wide analysis of the same tumours (see Fig. 5a in Aitken et al. 2025 (Aitken et al. 2025)), where the first two principal components capture strain and tumour status, with CAROLI most divergent.

**Figure 5.**
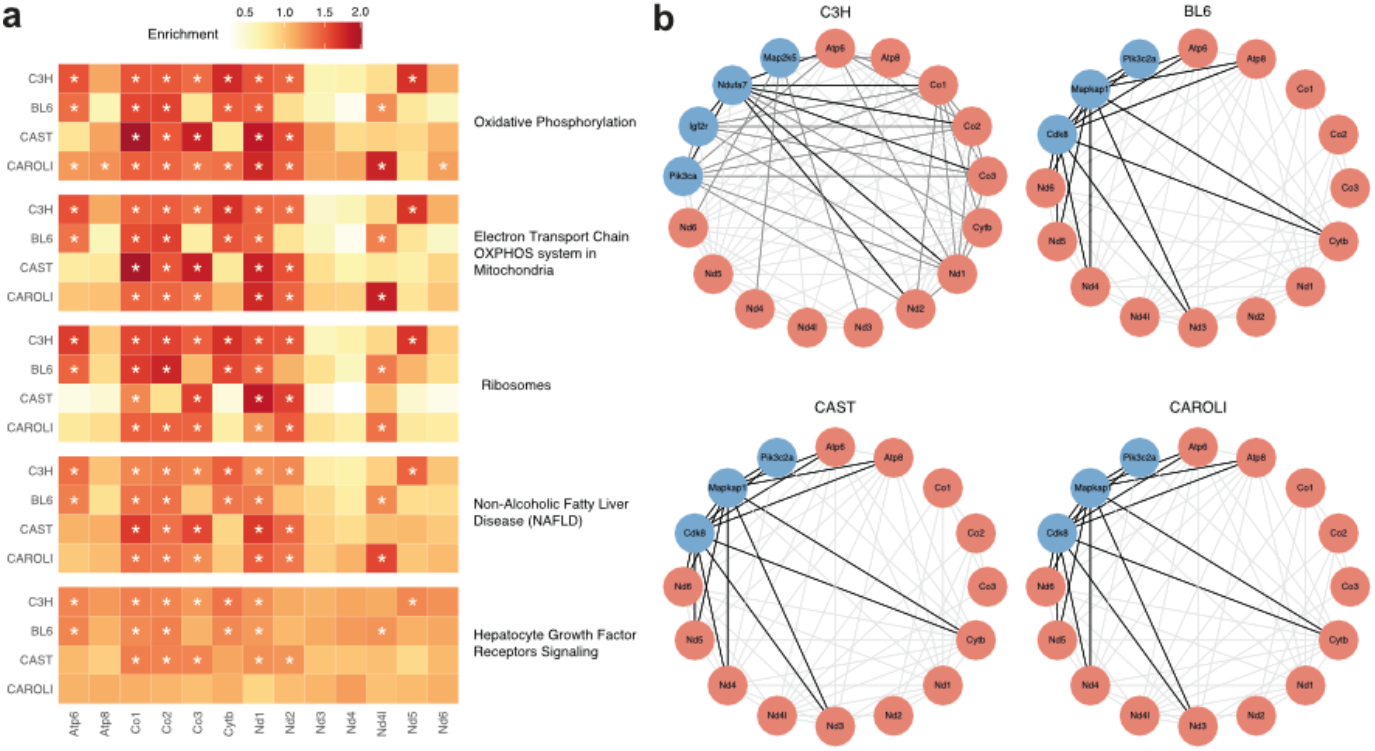
Co-expression networks link mitochondrial genes with nuclear genes. **(a)** Gene set enrichment analysis was performed using nuclear and mitochondrial genes identified from weighted gene co-expression network analysis (Methods). The heatmap shows the enrichment for each gene, per strain, for the top five ranked pathways. Enrichment levels are indicated by the colour gradient, with darker shades representing higher enrichment. Statistically significant enrichments are marked with an asterisk (* = p<0.05). (**b**) mtDNA-centric co-expression networks of C3H, BL6, CAST, and CAROLI. The networks illustrate the co-expression connections (grey/black edges) between mitochondrial genes (red nodes) and top nuclear genes (blue nodes); black edges indicating links that are at least 2-fold stronger than grey edges.

Here we have demonstrated that both genetic background and acquired somatic driver mutations in the nuclear genome impact mtDNA expression. The reduction in global mitochondrial gene expression in C3H, CAST, and CAROLI tumours is predominantly driven by reduced expression of the NADH complex genes, as well as *Atp6* and *Cytb*. The reduction of mtDNA expression observed in three of our four study strains has also been observed in human tumour samples, including liver cancer (Qiao et al. 2017).

### Co-expression mtDNA genes with MAPK pathway genes across strains

We studied the functional interactions between mtDNA and nuclear genes using weighted gene correlation network analysis (WGCNA; Methods) to calculate co-expression networks for the tumours from each of the four strains. This resulted in a list of co-expressed nuclear genes for each mtDNA gene. We then performed a gene set enrichment analysis (GSEA) for each strain, ranked by the strength of overall co-expression of each mtDNA gene (**Fig. 5a**). As expected, the nuclear genes most enriched for co-expression with mtDNA genes are found in pathways related to mitochondrial and liver function; for example, oxidative phosphorylation and electron transport chain in the OXPHOS system in mitochondria). In particular, one of the most enriched gene sets is the non-alcoholic fatty liver disease pathway, indicating that the mtDNA genes in our tumour samples are co-expressed with genes implicated in the pathogenesis of chronic liver disease.

We also generated co-expression networks for each of the strains (**Fig. 5b**) to identify nuclear genes that are strongly co-expressed with specific mitochondrial genes. All four strains show a strong connection between mitochondrial genes and nuclear genes found in the MAPK pathway: mitogen-activated protein kinase kinase 5 (*Map2k5*) for C3H, and mitogen-activated protein kinase associated protein 1 (*Mapkap1*) for BL6, CAST and CAROLI. In addition, the latter three strains also show co-expression with phosphatidylinositol-4-phosphate 3-kinase catalytic subunit type 2 alpha (*Pik3c2a*) and cyclin dependent kinase 8 (*Cdk8*). This analysis identifies a correlation between tumour mitochondrial gene expression and genes that play a role in cancer development. Understanding these mechanisms could help identify therapeutic strategies that target the mtDNA.

## Discussion

The rising burden of hepatocellular carcinoma (HCC) reflects changing environmental exposures and the increasing prevalence of chronic liver disease across diverse populations. Since reprogramming of cellular metabolism is a core hallmark of cancer (Hanahan and Weinberg 2011) that is characteristic of HCC, in this study we used a model system of chemical carcinogenesis to induce liver tumours in four genetically distinct mouse strains, and performed high resolution molecular mitochondrial profiling. This approach addresses the critical need to understand the interactions between carcinogenic exposures, mitochondrial function, and genetic variation. This model system also enabled us to control for the genetic variability and uncertain environmental exposures that characterise human studies.

We developed a novel, high-confidence heteroplasmy detection workflow that led to the identification of two types of mutations. First, “strain-recurrent” heteroplasmies, which are present in over 10% of tumour samples, appear to result from low-level germline heteroplasmies enriched in tumour cells but lost in normal tissues, likely due to random drift (He et al. 2010; Ju et al. 2014). Second, somatically acquired heteroplasmies, which accumulate during tumour clonal expansion (Ju et al. 2014), represent 51% of the identified heteroplasmies and are predominantly C>T and T>C substitutions. This mutational spectrum aligns with those seen in human cancers (Ju et al. 2014; Yuan et al. 2020) and likely arises from replication-coupled errors linked to bidirectional initiation of mtDNA genome replication (Ju et al. 2014).

Tumour samples exhibit significantly higher mtDNA mutational burdens compared to normal tissues, coupled with a striking depletion of mitochondrial content and gene expression. These appear to be conserved features of liver tumour development, rather than DEN treatment or age, since these differences are reproducible both within and across strains. These results corroborate observations from human studies (Yuan et al. 2020; Ju et al. 2014; Reznik et al. 2016; Zhang et al. 2019b), including an association between WGD and mtDNA content (Kim et al. 2024).

We present evidence that particular driver gene variants in the nuclear genome are linked to different levels of mitochondrial biogenesis. In addition, genome-wide co-expression analysis identified strong correlations between mitochondrial protein-coding genes and many nuclear genes, including an enrichment for MAPK pathway genes underscoring the crosstalk between nuclear and mitochondrial genes in tumourigenesis. Furthermore, these dramatic alterations to the mitochondrial genome and expression patterns are evident even in the earliest stage of liver tumours (dysplastic nodules) we have studied.

C3H mice, the strain most susceptible to cancer, exhibit the greatest depletion of mitochondrial expression. In contrast, CAROLI mice, the most cancer resistant strain, display a lower heteroplasmy burden and stronger purifying selection. The magnitude of differential expression also broadly correlates with tumour susceptibility with C3H showing the largest change and CAROLI the smallest. Previously, mitochondrial content has been associated with clinical features, such as the degree of hepatic steatosis (intracellular lipid accumulation) (Kamfar et al. 2016), and here we find that differences in mtDNA content and expression are associated with specific nuclear driver genes. For example, in CAST mice, *Egfr* driven tumours have the highest mtDNA content (**Fig. 3d**), which is also reflected in the mitochondrial expression (**Fig. 4b**).

Although the study’s design did not permit investigation of age-related mitochondrial changes, the absence of significant age effects across strains, despite varying tumour latencies, is intriguing. These findings suggest that certain mitochondrial features, such as expression changes, may represent universal responses to liver tumourigenesis, while others are strain-specific, likely influenced by evolutionary distance or nuclear genotype. Such trends are also seen in the nuclear genome, where we have identified that interactions between germline variation and the environment are influential in tumour genome evolution (Aitken et al. 2025).

In summary, our study extends understanding of the mitochondrial contributions to tumourigenesis and underscores the importance of considering genetic background in cancer research. The interplay between nuclear and mitochondrial genomes remains a promising avenue for future investigation, with potential for identifying novel therapeutic targets.

## Methods

### Tumourigenesis experiments

Liver tumours were induced, isolated, and processed as described previously (Connor et al. 2018; Aitken et al. 2020, 2025). Briefly, 15-day-old male mice were treated with a single intraperitoneal dose of diethylnitrosamine (DEN), and tumours isolated at fixed time points after DEN treatment (C3H 25 weeks, BL6 36 weeks, CAST 38 weeks, CAROLI 78 weeks). Tumours bisected, with one half for histopathology and the other half processed for DNA and RNA extraction and whole genome (WGS) and total transcriptome sequencing (RNA-seq). Additional normal liver samples were processed for control experiments (**Table S1**).

Whole genome sequencing of normal liver and liver tumours are available in the European Nucleotide Archive (ENA) under accession PRJEB37808 (C3H and CAST) (Aitken et al. 2020) and PRJEB15138 (BL6 and CAROLI) (Aitken et al. 2025). Total transcriptome sequencing of liver tissue and tumours is available at ArrayExpress (accession pending) (Aitken et al. 2025). Samples and associated metadata are listed in **Table S1**.

### Novel *de novo* mitochondrial genome assembly and annotation for C3H, CAST, and CAROLI mice

The strains (reference genome assembly) used in this study are *Mus musculus musculus* C57BL/6J (GRCm39), *Mus musculus musculus* C3H/HeOuJ (C3H/HeJ), *Mus musculus castaneus* CAST/EiJ (CAST/EiJ) and *Mus caroli* CAROLI/EiJ (CAROLI_EiJ) from Ensembl v.91 (Howe et al. 2021).

BL6 was analysed using the NC_005089.1 C57BL/6J reference mitochondrial genome accessed from NCBI. The C3H, CAST and CAROLI mitochondrial genomes were assembled *de novo* using NOVOplasty v. 3.8.1 (Dierckxsens et al. 2017, 2020) and have been deposited in GenBank (PV177089, PV231059, PV231060). For this, WGS of all normal liver tissue samples were used for each strain (C3H=4; CAST =7; CAROLI=5).

For C3H and CAST, NOVOplasty was run on all the WGS libraries with default parameters using the *Cytb* gene sequence of NC_005089.1 as seed. CAROLI exhibits evolutionary divergence of *Cytb* gene and thus required a two-step process: first, the NC_005089.1 *Cytb* gene was used a as seed to create a preliminary assembly from which the longest conserved sequence between the preliminary assembly and NC_005089.1 was used as a seed for NOVOplasty to create the final assembly. All libraries for all samples returned circularised assemblies. For all strains, the assemblies for the various libraries were essentially identical (assemblies from two C3H libraries had a Y at one position that was an A in all others) and used to create strain-specific consensus circularised reference mitochondrial genomes assemblies for each strain. Gene annotation of the three mitochondrial genomes were performed using MITOS (Bernt et al. 2013). Start and stop positions for all genes were manually curated. The genes identified for the three strains are identical and matched BL6 gene annotation (**Fig. 1**).

To compare sequence similarity of the four mitochondrial genomes, multiple sequence alignment was performed using Clustal Omega (Madeira et al. 2024).

See **Table S2** for the final assembly and annotation.

### Comparative mitochondrial genome analysis

For each strain, we aligned our *de novo* assembly and the corresponding sequences retrieved from GenBank using Clustal Omega with “sequence type” set to DNA. A summary of all sequences included in the comparative analysis is provided in **Table 1**.

### Heteroplasmy detection and annotation

We mapped the trimmed WGS libraries using the bwa-mem function from BWA (v.0.7.17) (Li and Durbin 2009), using the mitochondrial genomes as reference. Valid mapped pairs to the mitochondrial genome were then extracted using the view function from Samtools (v.1.9) (Li et al. 2009). The remaining reads were then re-mapped to the full genome (nuclear and mitochondrial) using bwa-mem. Valid mapped pairs to the mitochondrial genome were extracted a second time. This approach allows potential NUclear Mitochondrial Sequences (NUMTs) to be filtered out, which otherwise tend to misalign to the mitochondrial genomes because of their high identity levels with the mitochondrial genome (Zhang et al. 2019b). Duplicates were removed using the MarkDuplicates function from Picard Tools (v.2.21.7) ([CSL STYLE ERROR: reference with no printed form.]).

For BL6, GRCm39 was used as the nuclear genome assembly to address a BL6 specific issue caused by a problematic NUMTs (Numts_002 in (Calabrese et al. 2012)) sequence located on 1: 24,650,616-24,655,269 (4,654bp long) that is 99.94% identical to MT:6,390-11,042 (4,653bp long). Although this region was improved in GRCm39, optimal mapping quality on the BL6 mitochondrial assembly required masking of the region on the nuclear genome. The masked sequence is within a non-coding region of the BL6 nuclear genome and is not conserved in the other strains. We are confident that masking this sequence did not have an impact on the heteroplasmy detection quality.

The mpileup function from Samtools was then used to generate pileup files from the BAM files with “-E -f” parameters to optimise the accuracy of the heteroplasmy detection (Wang et al. 2020). Heteroplasmy were detected using the mpileup2snp function of VarScan (v.2.4.4) (Koboldt et al. 2012). Finally, heteroplasmies were classified as synonymous, missense, loss of function, tRNA, rRNA, and non-coding using snpEff (v.5.1.2) using default parameters (Cingolani et al. 2012).

The mitochondrial genome is circular and must be arbitrarily cut to meet the input requirements of mapping alignment software. To mitigate biases caused by this linearisation (apparent low read coverage at the “ends”), we created a shifted version of each of the mitochondrial genomes with the starting position shifted by 8,150bp (**Fig. 1, Fig. S3b**). We re-applied the mapping approach described above using the shifted version of the mitochondrial genomes. The final mutation set was obtained by taking the mutation from the reference genome mapping from position MT:1,100-15,200 and the mutation from the shifted genome mapping from position MT:1-1,099 and MT:15,201-end of the genome. Re-mapping the libraries on a shifted mitochondrial genome ensures a deep coverage at every position of the mitochondrial genome.

A total of 3,382 heteroplasmies were initially detected, corresponding to 1,717 unique mitochondrial genomic positions.

To account for hypermutated samples (**Table S1, Fig. S6**), we first excluded any sample containing more than 50 heteroplasmies, which led to the removal of three samples (two from CAST and one from C3H). To further refine the filtering, we excluded samples with a heteroplasmy count exceeding three standard deviations from the strain-specific mean, resulting in the additional removal of four samples (two from CAST and two from CAROLI). After this filtering step, 2,232 heteroplasmies remained, covering 1,011 mitochondrial genomic positions (**Fig. 1**).

We further classified the remaining heteroplasmies into two categories: strain-recurrent and somatic. A heteroplasmy was defined as “strain-recurrent” if it was present in more than 10% of the strain’s samples (or in more than 10 samples for C3H). After filtering out strain-recurrent heteroplasmies, 1,138 heteroplasmies remained, corresponding to 975 unique mitochondrial genomic positions. See **Table S4** for the list of all identified heteroplasmies and associated annotations.

See GitLab (https://gitlab.com/maelle.daunesse/lce-mtdna.git) for complete snakemake pipeline.

### Heteroplasmy analysis

Sequence-composition adjusted mutation rates were calculated as defined previously (Aitken et al. 2020) with modification to consider the mitochondrial genome:

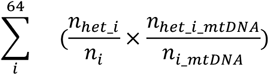

Where *n*_*het_i*_ corresponds to the number of the trinucleotide *i* with a heteroplasmy in the gene of interest, *n*_*i*_ corresponds to the number of the trinucleotide *i* in the gene of interest, *n*_*het_i*_*mtDNA*_ corresponds to the number of the trinucleotide *i* with a heteroplasmy in the mitochondrial genome and *n*_*i*_*mtDNA*_ corresponds to the number of the trinucleotide *i* in the mitochondrial genome.

dN/dS ratios were calculated for each gene using the dNdScv R package (v1.0) (Martincorena et al. 2017), which estimates the ratio of nonsynonymous to synonymous substitution rates while accounting for mutation rate variation and sequence composition biases. dNdScv employs a trinucleotide mutation model to correct for context-dependent mutational effects and was run using the “Vertebrate Mitochondrial Code” (`numcode = 2`) to appropriately model mitochondrial coding sequences. dN/dS scores are listed in **Table S3**.

SIFT scores were calculated for each missense annotated heteroplasmy using SIFT (v.6.2.1) (Kumar et al. 2009), with alignments calculated by NCBI BLAST+ (v.2.4.0) (Altschul et al. 1997) and UniRef90 (release 2025_01) as the reference protein database. SIFT scores are listed in **Table S4**.

See GitLab (https://gitlab.com/maelle.daunesse/lce-mtdna.git) for complete Rmarkdown notebooks.

### Mitochondrial content calculation

We mapped the trimmed reads to the full genome (mtDNA + nuclear DNA) and duplicates were removed using the MarkDuplicates function from Picard. Finally, we used the idxstat function from Samtools to obtain the number of reads mapped to each chromosome.

Mitochondrial content was estimated using deduplicated libraries by calculating the fraction of mitochondrial reads in each of the WGS libraries (**Table S5**). We applied a formula that accounts for both cellularity and ploidy, as adapted from Reznik et al. 2016 (Reznik et al. 2016), which originally used a tumour purity metric (here replaced with cellularity). The cellularity of a tumour corresponds to the estimated fraction of sequenced cells that represent the dominant clonal expansion in the tumour (between 0 and 1); for normal samples, the cellularity was imputed as 1).

The mitochondrial content was calculated as follows:

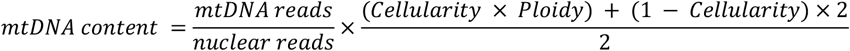

This adjustment ensures the mitochondrial content reflects the cellular context, accounting for tumour heterogeneity and variations in ploidy.

### Mitochondrial gene expression analysis

We mapped the trimmed RNA-seq libraries generated from normal liver tissue (15-day old and adult) and tumour samples using STAR (v.2.7) (Dobin et al. 2013), using the full genomes described above (nuclear and mitochondrial) as a reference. Read counts per gene were obtained using the featurecounts function from Subread (v.2.0.3) (Liao et al. 2014). Only the 14,779 genes that are orthologous in the four strains were kept for the next steps (**Table S6**).

Read counts from each library were then normalised using DEseq2 (v.1.38.2) (Love et al. 2014). We performed differential expression using normal adult and DN tumour samples with DESeq2 accounting for the strain, diagnosis, and cellularity in the design (**Table S6**). The PCA shown in **Fig. 4d** was obtained after variance stabilising transformations (VST). The 15-day old and adult normal samples cluster together (**Fig. S10)**.

### Co-expression of mitochondrial genes analysis

We performed a co-expression analysis using the WGCNA R package (v.1.71) (Langfelder and Horvath 2008). For each strain, a weighted gene co-expression network was built using both mitochondrial and nuclear genes using a soft thresholding power (β) of 5. The resulting network assigns each pair of genes a weight ranging from 0 (low) to 1 (high) that corresponds to the connection strength between the two genes.

To determine the pathways with which the mitochondrial genes are co-expressed, we used a gene-set enrichment analysis (GSEA) approach. The GSEA analysis was performed using the co-expression weights as input to rank the genes, employing the fgsea R package (v.1.24.0) (Korotkevich et al. 2021). Finally, we built a mtDNA-centric network using the 100 most connected nuclear genes to all the mitochondrial ones.

## Computational analysis environment and code availability

The analysis pipeline including snakemake pipelines to process the data, subsequent scripts and R notebook to generate the figures can be obtained without restriction from the repository https://gitlab.com/maelle.daunesse/lce-mtdna.git.

## Data availability

Sequencing data in this study are described in Aitken et al. 2020 (Aitken et al. 2020) and Aitken et al. 2025 (Aitken et al. 2025). Whole genome sequencing of normal tissues and and liver tumours are available in the European Nucleotide Archive (ENA) under accession PRJEB37808 (Aitken et al. 2020) and PRJEB15138 (Aitken et al. 2020, 2025). Total transcriptome sequencing of liver tissue and tumours is available at ArrayExpress (accession pending) (Aitken et al. 2025). Whole genome sequencing alignment for reference and shift mapping of the mitochondrial genomes of liver tissue and tumours are available in NCBI under accession PRJNA1225162. The mitochondrial genomes assembled for C3H, CAST, and CAROLI in this study have been deposited in GenBank (PV177089, PV231059, PV231060).

## Ethics

Animal experimentation was carried out in accordance with the Animals (Scientific Procedures) Act 1986 (United Kingdom) and with the approval of the Cancer Research UK Cambridge Institute Animal Welfare and Ethical Review Body (AWERB). Full details of the experimental procedures leading to the data generation are provided in Aitken et al. 2020 (Aitken et al. 2020) and Aitken et al. 2025 (Aitken et al. 2025).

## Acknowledgements and funding

We thank Andrew Yates (EMBL-EBI) for assistance with data management, and members of the Aitken Lab (H. Ahmed Nur, C. Arnedo-Pac, L. Cocker, E. Reilly) for comments on the manuscript.

This work was supported by European Molecular Biology Laboratory core funding, Cancer Research UK Cambridge Institute core award (20412), MRC Human Genetics Unit core funding programme grants (MC_UU_00007/11, MC_UU_00007/16, MC_UU_00035/1, MC_UU_00035/2), and MRC Toxicology Unit core funding programme grants (RG94521 and MC_PC_24012). IRB Barcelona is a recipient of a Severo Ochoa Centre of Excellence Award from the Spanish Ministry of Science, Innovation and Universities (MICINN, Government of Spain) and is supported by CERCA (Generalitat de Catalunya). Support was also provided from specific research grants: Cancer Research UK strategic award (22398); Wellcome Trust (WT108749/Z/15/Z, WT202878/B/16/Z, 202878/Z/16/Z); PID2021-126568OB-I00 (CHEMOHEALTH) project, funded by the Spanish Ministry of Science (MCIN), AEI /10.13039/501100011033; ERC (615584, 788937); Helmholtz Society (DKFZ abteiling B270). J.C. was supported by a Wellcome Trust PhD Training Fellowship for Clinicians (WT223088/Z/21/Z) as part of the Edinburgh Clinical Academic Track (ECAT) programme. M.D. and E.K. were supported by the EMBL International PhD Programme. V.S. received an EIPOD fellowship. S.J.A. received a Wellcome Trust PhD Training Fellowship for Clinicians (WT106563/Z/14/Z and WT106563/Z/14/A), National Institute for Health Research (NIHR) Clinical Lectureship, and CRUK Clinician Scientist Fellowship (RCCCSF-May23/100001).

## Author contributions

M.D., S.J.A., and P.F. conceptualised the project. D.T.O. supervised experimental data collection. M.D. and P.F. performed primary genomic and sequence data analysis. V.S., E.K. J.C., and C.J.A. provided computational support. N.L.B., C.S., M.S.T., and M.R. advised on specific analysis components. S.J.A. and P.F. supervised the work. M.D., S.J.A., and P.F. wrote the paper. All authors had the opportunity to edit the manuscript. All authors approved the final manuscript.

The Liver Cancer Evolution Consortium members are:

Sarah J. Aitken, Stuart Aitken, Craig J. Anderson, Claudia Arnedo-Pac, John Connelly, Frances Connor, Maëlle Daunesse, Ruben M. Drews, Ailith Ewing, Christine Feig, Paul Flicek, Paul A. Ginno, Vera B. Kaiser, Elissavet Kentepozidou, Erika López-Arribillaga, Núria López-Bigas, Juliet Luft, Margus Lukk, Duncan T. Odom, Oriol Pich, Tim F. Rayner, Colin A. Semple, Inés Sentís, Vasavi Sundaram, Lana Talmane, Martin S. Taylor & Jan C. Verberg

