## Supplemental Figures S1-S10 for "Germline and somatic variation influence nuclear-mitochondrial crosstalk in tumourigenesis"

### Supplementary Figures

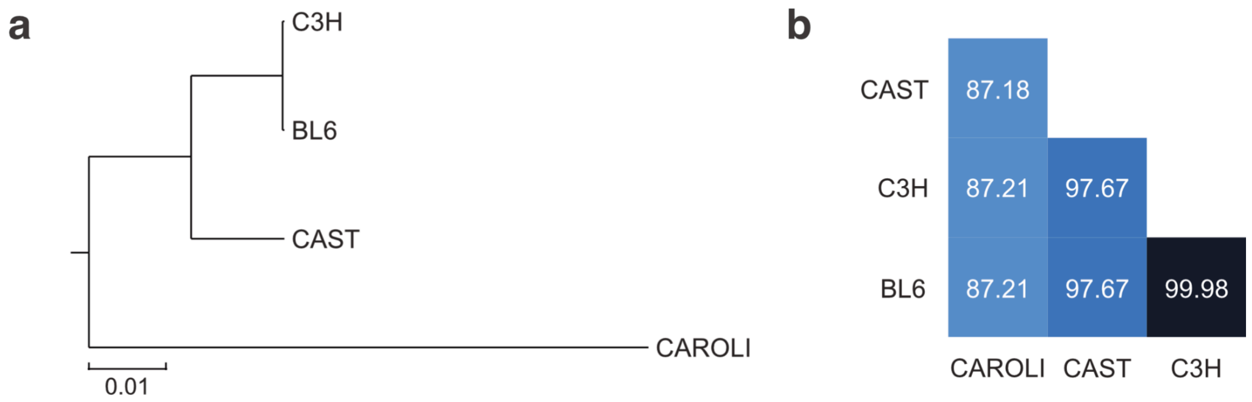


**Figure S1 | *De novo* assembly metrics of C3H, CAST and CAROLI mitochondrial genomes.**

**(a)** Phylogenetic tree of the four mitochondrial genomes obtained through Clustal Omega multiple sequence alignment. The scale bar corresponds to the number of substitutions per site, i.e. the average number of nucleotide or amino acid changes that occur at each position across all sequences in the alignment. **(b)** Pairwise comparison of each of the four mitochondrial genomes (percentage similarity shown) identifies that sequence divergence follows evolutionary distance.

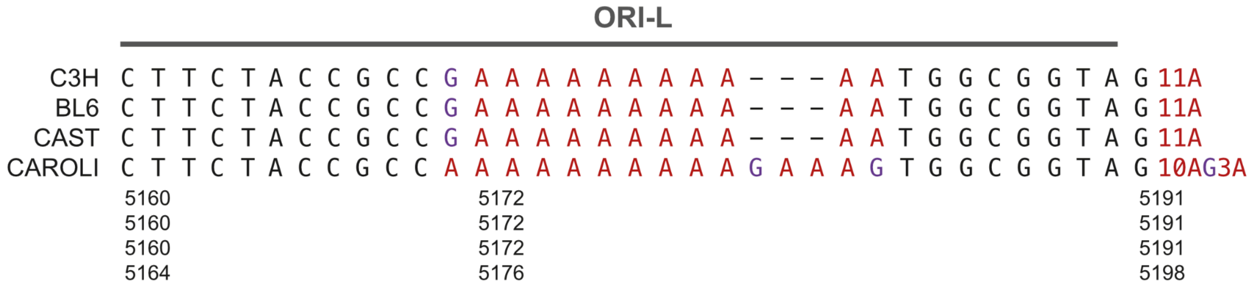


**Figure S2 | The origin of replication of the light strand (ORI-L) is not conserved in CAROLI.**

The ORI-L sequence is shown for each of the four strains, highlighting conserved and variable regions. In C3H, BL6, and CAST, the ORI-L sequence is fully conserved, including an 11-adenine repeat (red), whereas this region has expanded to 10AG3A in CAROLI. The mitochondrial genomic coordinates are shown underneath for each of the strains.

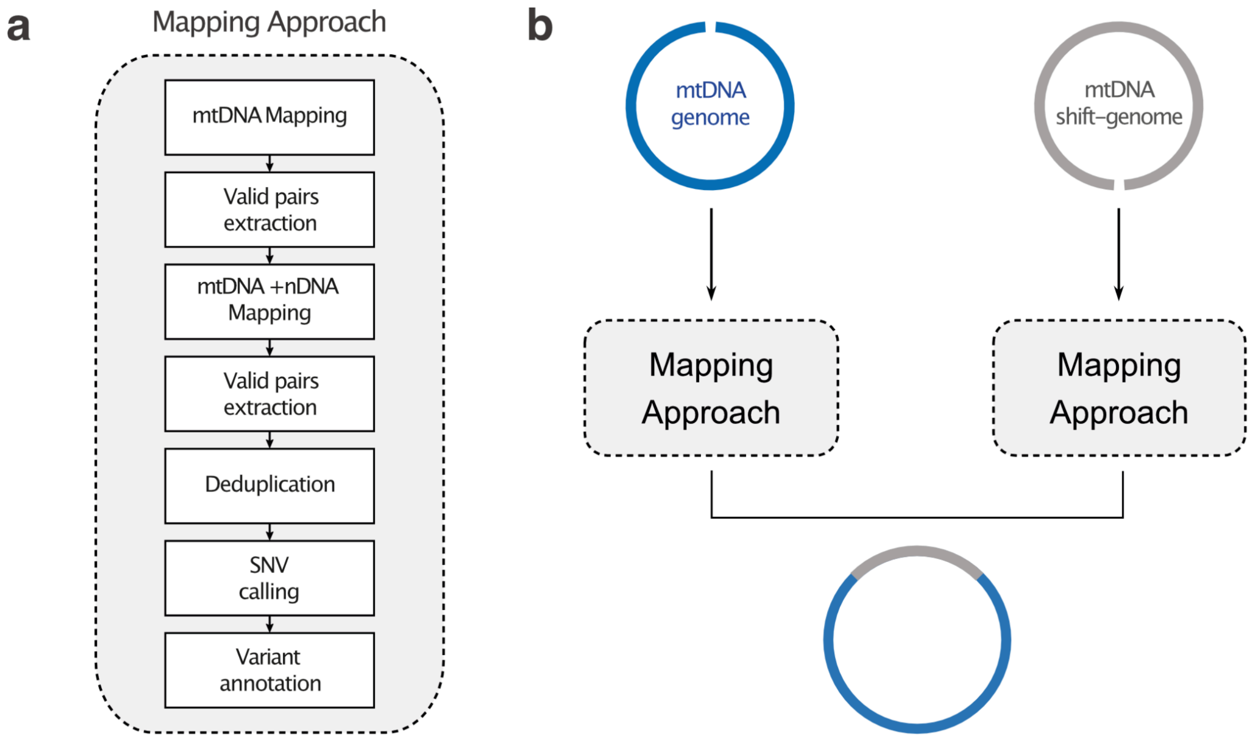


**Figure S3 | Heteroplasmy detection method overview.**

**(a)** Detailed mapping approach designed to minimise the misalignment of nuclear mitochondrial reads (NUMTs). The process includes mtDNA mapping, extraction of valid pairs, combined mitochondrial DNA (mtDNA) and nuclear DNA (nDNA) mapping, further extraction of valid pairs, deduplication, single nucleotide variant (SNV) detection, and variant annotation. **(b)** Due to the circular nature of mtDNA, the mapping approach (shown in **a**) is applied using both the reference mtDNA genome and a shifted version of the mtDNA genome. This ensures comprehensive coverage, particularly at the ends of the reference mtDNA.

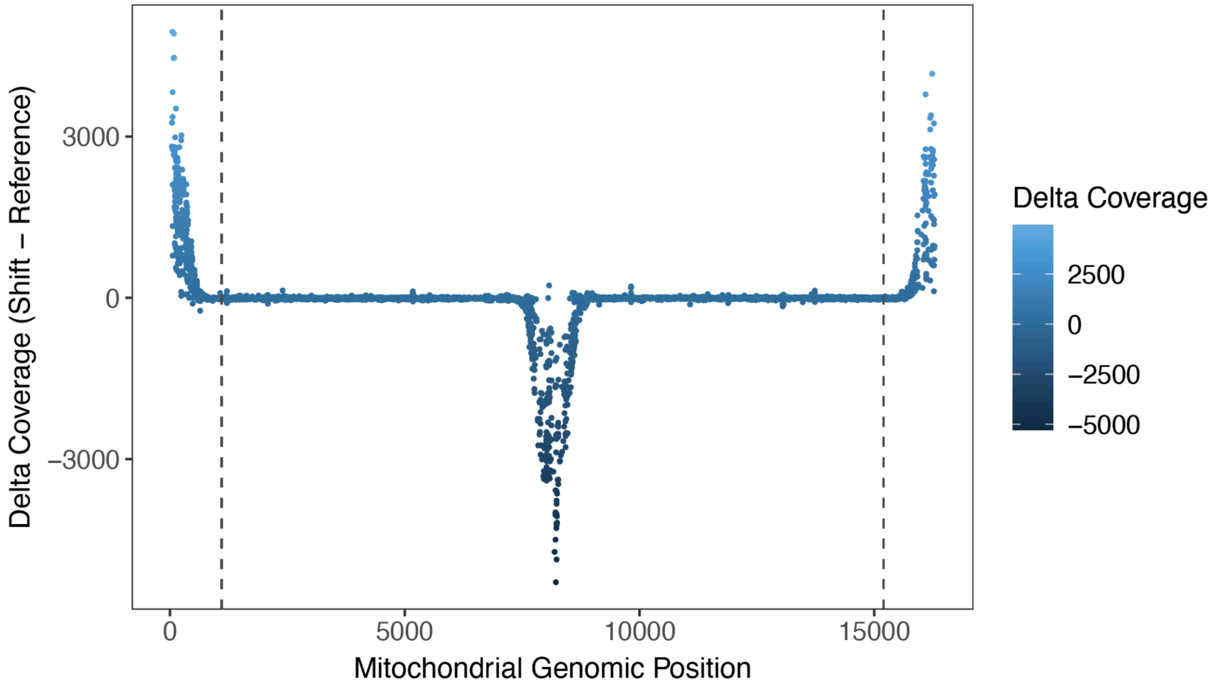


**Figure S4 | The effect of shifted mapping on mitochondrial read coverage.**

Differences in mitochondrial read coverage between classic and shifted mapping approaches. Sequencing coverage of the mitochondrial genome across all strains was quantified separately for the classic mapping approach and the shifted mapping approach (**Fig. S3**), and the delta coverage is plotted. Positive values on the y-axis indicate higher coverage in the shifted mapping, while negative values indicate higher coverage in the classic mapping. The differences correspond to the artificial cut point of the circular mitochondrial genome, and corresponding lower coverage, in each approach (plotted in **Fig. 1**).

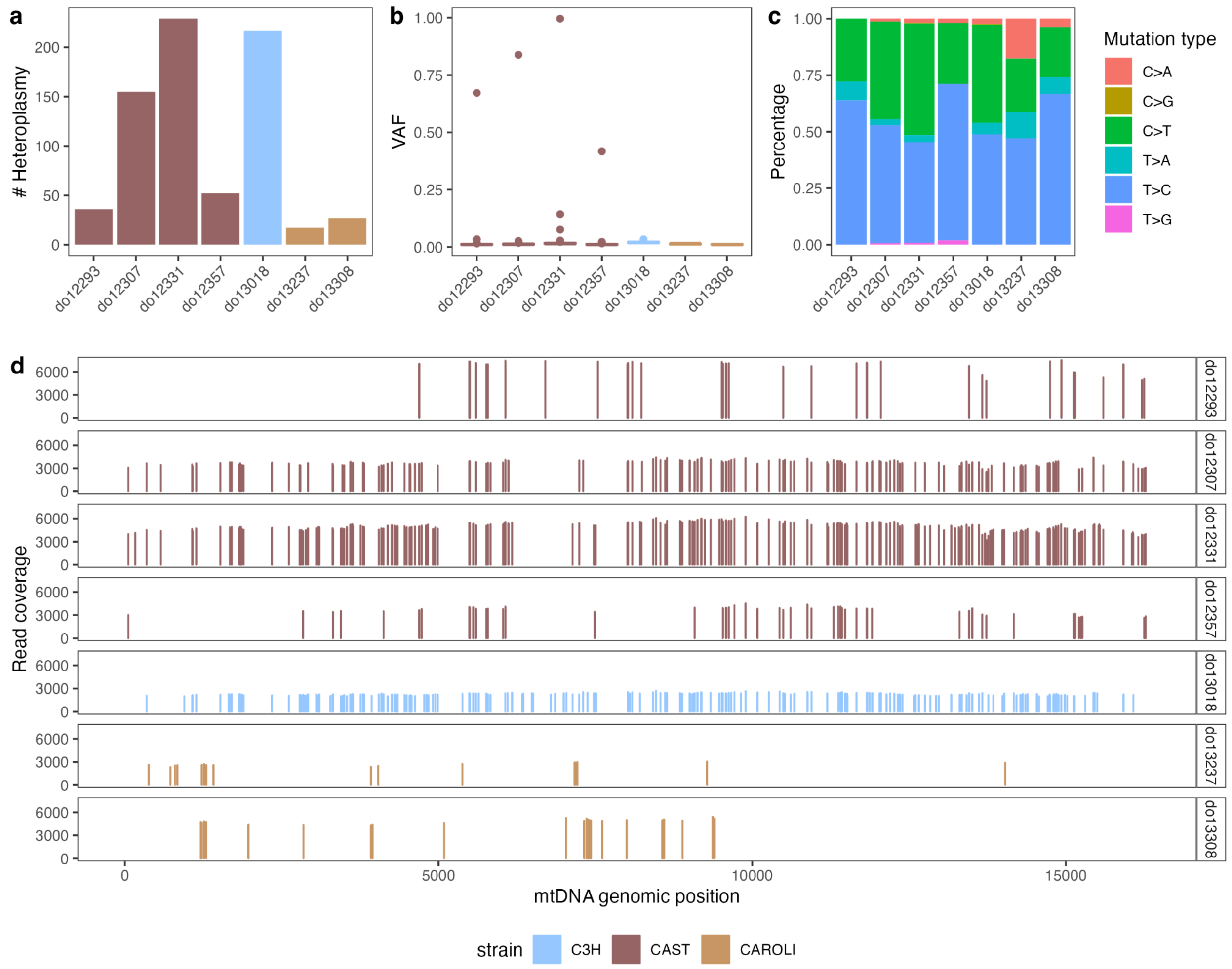


### Figure S5 | Hypermutated sample profile

(**a**) Number of heteroplasmic mutations detected mtDNA for each sample (n=7). (**b**) Variant allele frequency (VAF) distribution of heteroplasmic mutations across samples. (**c**) Proportions of mutation types among heteroplasmic sites in each sample. (**d**) Read coverage of each detected heteroplasmy. Strains are colour-coded as follows: C3H (blue), CAST (brown), and CAROLI (orange).

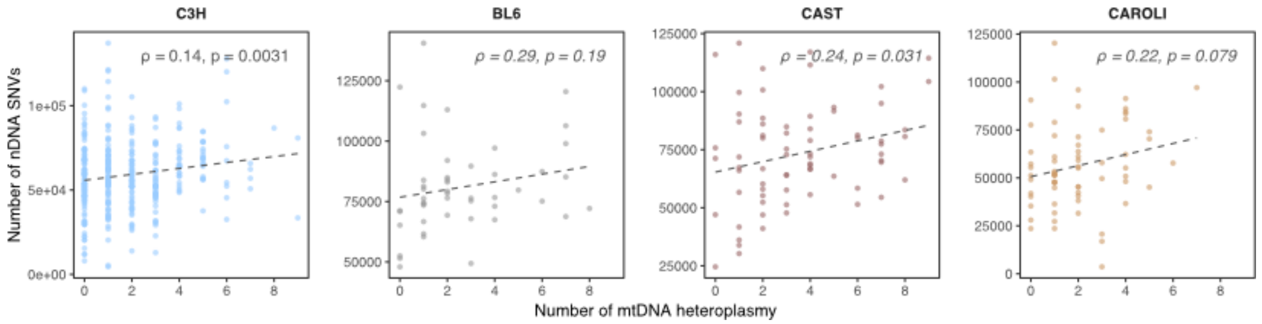


***Figure S6 | Positive correlation between heteroplasmies and nuclear somatic mutations.***

In tumours, the number of mitochondrial heteroplasmies (x axis) correlates positively with the number of nuclear DNA (nDNA) single nucleotide variants (SNVs) (y-axis). Spearman correlation coefficients (ρ) are plotted and corresponding p-values (p) are indicated for each strain.

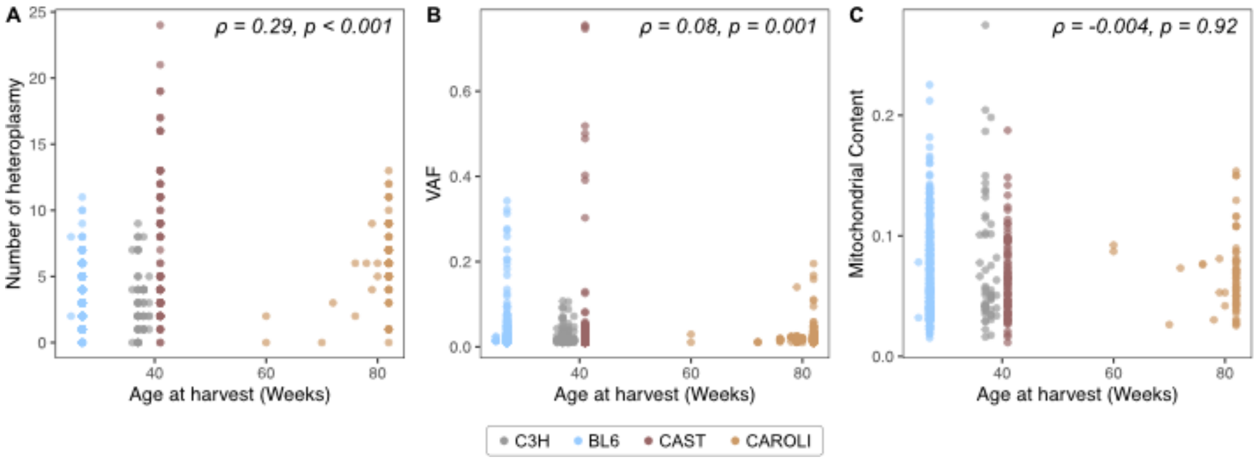


**Figure S7 | Mouse age is only weakly correlated with mitochondrial features.**

**(A)** The number of heteroplasmies (y axis) per tumour is not correlated with the age at the time of tumour isolation. **(B)** The variant allele frequency (VAF) of heteroplasmies is not correlated with age. **(C)** Mitochondrial content does not correlate with age. Notably, the experimental design confounds this analysis due to fixed time points for tumour isolation per strain (Aitken et al. 2025). Colours indicate the mouse strain. Correlation coefficients are indicated for each strain. ρ = Spearman correlation coefficient, p = associated p-value.

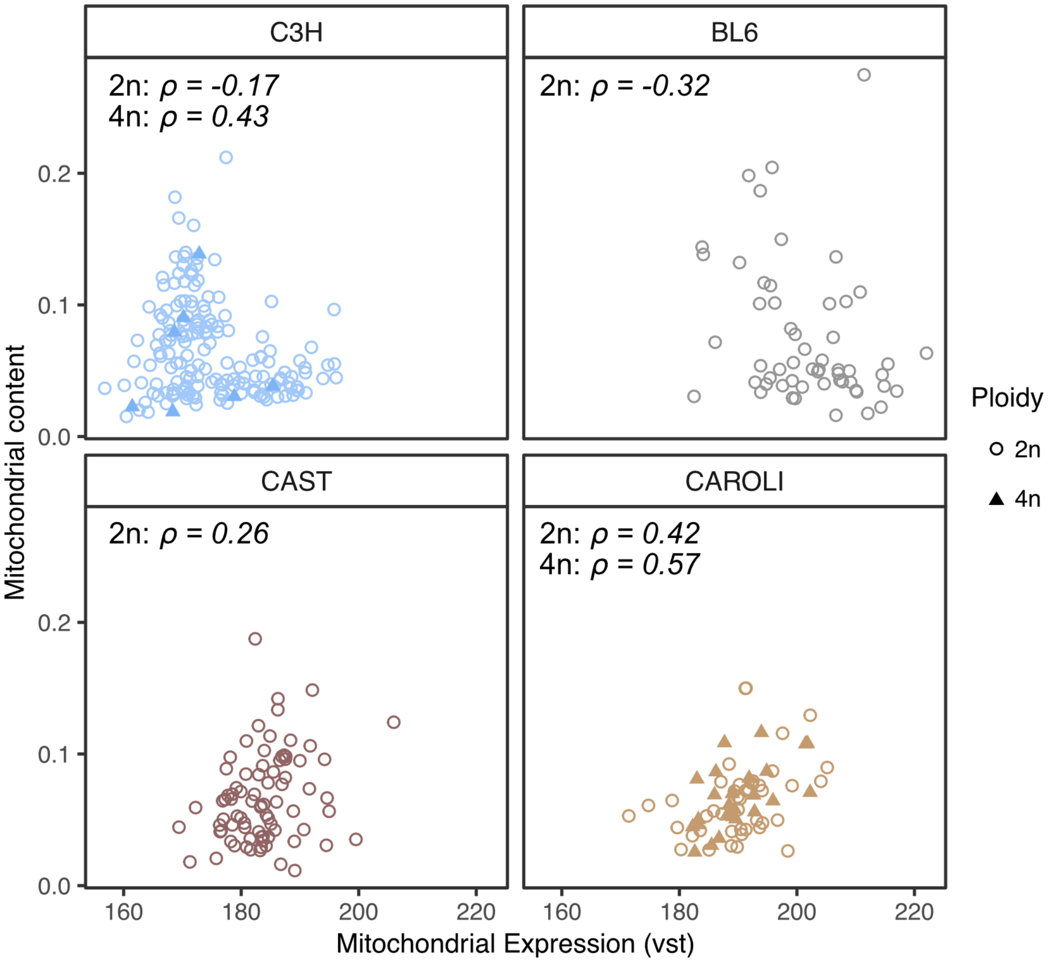


**Figure S8 | mtDNA content and tumour gene expression are correlated**

Spearman correlation (ρ) was calculated between the mtDNA content and expression for tumours samples (DEN-treated DN) with paired WGS and RNA-seq data. Tumours are grouped by ploidy, where 2n indicates diploid tumours and 4n corresponds to whole genome duplicated (WGD) tumours. mtDNA content and gene expression are moderately correlated in WGD tumours (found only in C3H and CAROLI) as well as diploid CAROLI tumours.

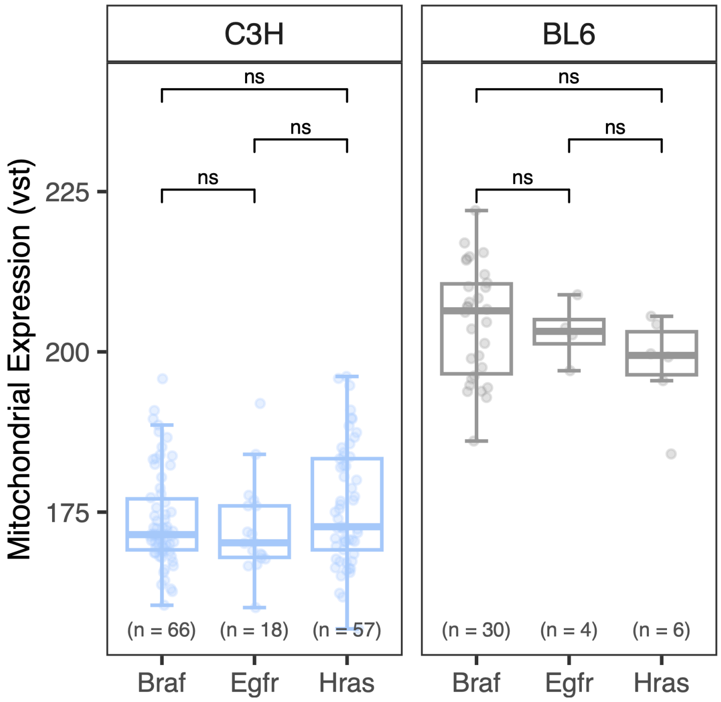


**Figure S9 | Mitochondrial expression is not correlated with driver genes in C3H or BL6.**

In contrast with CAST (**Fig. 4b**), C3H and BL6 present no association between mitochondrial expression and driver gene mutations. Wilcoxon test (ns = p>0.05).

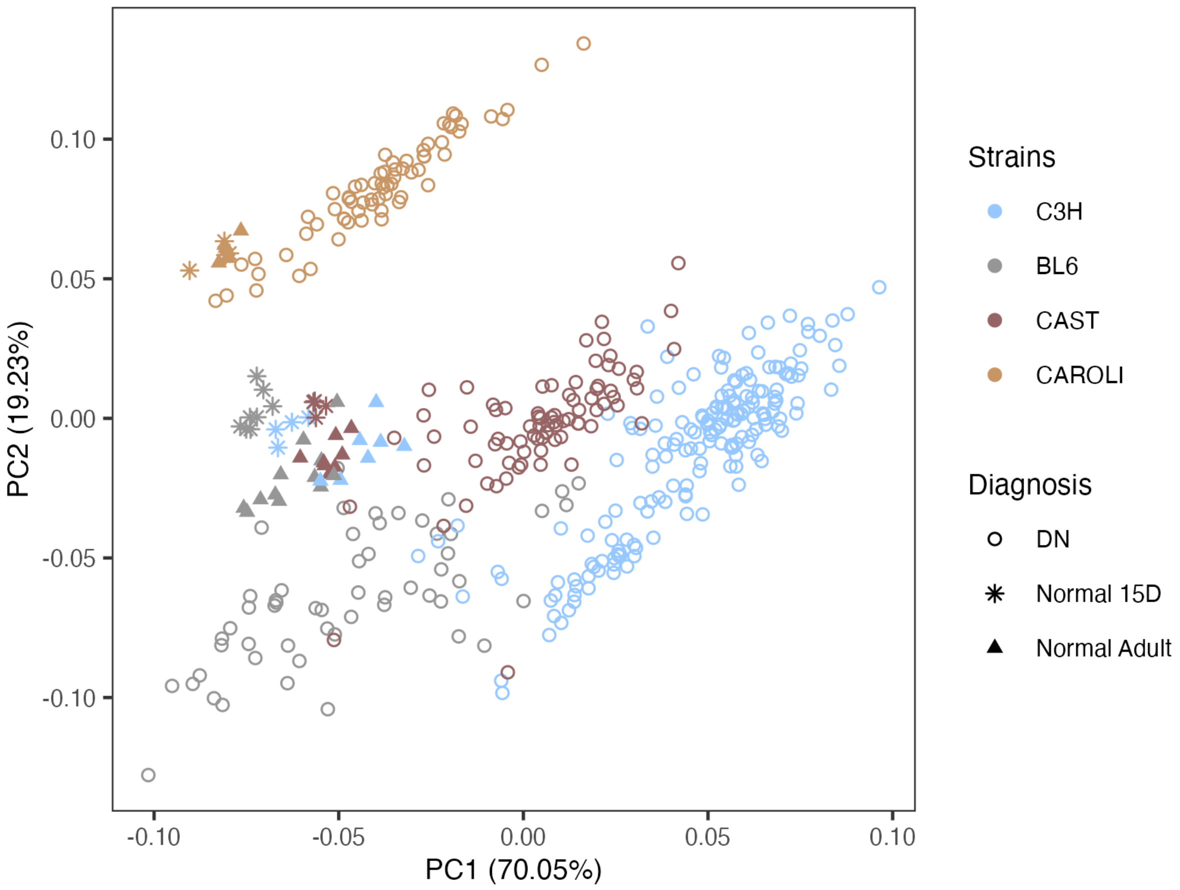


**Figure S10 | Principal component analysis with normal sub-group information**

Principal component analysis (PCA) of the variance stabilisation transformed (VST) mitochondrial read counts for 15-day-old normal (star), adult normal (triangle), and tumour samples (circle).
